# Blood vessels regulate primary motor neuronal pathfinding in zebrafish via exosome contained microRNA-22

**DOI:** 10.1101/2021.04.29.441918

**Authors:** Jiajing Sheng, Jie Gong, Yunwei Shi, Xin Wang, Dong Liu

## Abstract

A precise neuro-vascular communication is crucial to orchestrate directional migration and patterning of the complex vascular network and neural system. However, how blood vessels are involved in shaping the proper neuronal formation has not been fully understood. So far, limited studies have reported the discovery and functions of microRNAs (miRNAs) in guiding vascular and neural pathfinding. Currently, we showed that the deficiency of miRNA-22a, an endothelial-enriched miRNA, caused dramatic pathfinding defects both in intersegmental vessels (ISVs) and primary motor neurons (PMNs) in zebrafish embryos. Furthermore, we found the specific inhibition of miR-22a in ECs resulted in the patterning defects of both ISVs and PMNs. However, neuronal block of miR-22a mainly led to the axonal defects of PMN. Then we demonstrated that endothelial miR-22a regulates PMNs axonal navigation via exosome pathway. Sema4c was identified as a potential target of miR-22a through transcriptomic analysis and *in silico* analysis. Furthermore, luciferase assay and EGFP sensor assay *in vivo* confirmed the binding of miR-22a with 3’-UTR of *sema4c*. In addition, Down-regulation of *sema4c* in the miR-22a morphants significantly neutralized the aberrant patterning of vascular and neural networks. Our study revealed that miR-22a acted as a dual guidance cue coordinating vascular and neuronal patterning and expanded the repertoire of guidance molecules, which might be of use therapeutically to guide vessels and nerves in the relevant diseases that affect both systems.

## Introduction

During vertebrate embryogenesis, the formation of an architectural vasculature and nerve network is essential to ensure proper functioning. Blood vessels and nerves are exquisitely organized in nearly parallel patterns throughout the body. However, the mechanisms that regulate the ECs and neurons to follow specific migratory tracts during development have not been answered sufficiently. Recent reports have demonstrated that vasculature and nerves possess common molecular factors to guide cell migration and pathfinding and neuro-vascular communication is crucial for the development of both systems. Increasing evidences have demonstrated that several axon guidance signals including *Robos*, *UNC58*, *Plexins* and *Neuropilins* regulate vessel pattern and vice versa [1]. Apart from nerves, vessels also produce axon-guiding signals and vascular patterning signals [2, 3]. In addition, proper neuronal axonal wiring in brain development also depends on the precise molecular regulation of neuro-vascular co-patterning [4]. Despite recent progresses in the view of neuron control vascular guidance, the understanding of vessel contributions to nerves navigation and neuro-vascular communication is remained to be fully unexplored.

Semaphorins, categorized into eight classes, are a large family of transmembrane or secreted proteins, which have been studied as axonal guiding cues for years and also found to participate in the processes of vascular development recently [5, 6]. For example, like in the nervous system, autocrine endothelial Sema3A regulate endothelial cell migration, vascular navigation and patterning by binding to NRP1/Plexin [7]. Sema5A is a pro-angiogenic semaphorin that was suggested to regulate blood vessels remodeling and hierarchical organization during embryo development [8]. Class 4 subfamily of semaphorins have been found to be closely related to immunity and inflammation [9]. However, Sema4A, initially found as an activator of T cell-mediated immunity, was identified to promote endothelial cell apoptosis by inhibiting tyrosine phosphorylation of VEGFR2 [10]. In addition, Sema4D was found to increase endothelial cell migration through mDIA1/Src signaling pathway [11]. Above results indicated that class 4 semaphorins may also participate in behavior of endothelial cell. The Sema4c has been shown to play diverse diversity functional roles in biological development, including neurogenesis [12], terminal myogenic differentiation [13] and epithelial mesenchymal transition (EMT) [14]. In addition, Sema4c and its receptor Plexin-B2 have been demonstrated to express either in nervous system and endothelial cells, but knowledge of their functions in vascular development is still limited [15].

miRNAs, a class of small non-coding RNAs, are key post-transcriptional regulators to inhibiting the translation of target mRNAs through binding at the 3’untranslated region (UTR). A miRNA often regulates a cluster of targets in a variety of biological processes. Of interest, more miRNAs have been reported to be widely distributed in nerves and vascular system [16]. Thus, miRNAs are important candidate regulators of neurons and vessels development. A number of miRNAs have been reported to make roles during angiogenesis. However, the role of miRNAs in neuro-vascular development has not been extensively explored.

Currently, our understanding on the association between axon guidance and vascular pattern is still superficial. Here, by taking advantages of the zebrafish as a model, we addressed these questions and identified miR-22a derived from endothelial cells make a dual role in vascular and neuronal guiding by targeting *sema4c*. Knockdown of miR-22a caused zebrafish ectopic network in both blood vessel and nerves. The defect of the vascular patterning was mimicked by endothelial-specific reduction of the miR-22a. Interestingly, the phenotype in nerves network was induced by both neuron-specific and EC-specific repression of miR-22a. We revealed that endothelial miR-22a regulates PMNs axonal navigation via exosome pathway. Furthermore, we identified *sema4c* as a direct target of miR-22a. These observations demonstrate that blood vessels mediate primary motor neuronal pathfinding in zebrafish via exosome contained miR-22.

## Materials and Methods

### Ethics statement

All animal experimentation was carried out in accordance with the NIH Guidelines for the care and use of laboratory animals (http://oacu.od.nih.gov/regs/index.htm) and ethically approved by the Administration Committee of Experimental Animals, Jiangsu Province, China (Approval ID: 20150305-029).

### Zebrafish husbandry and breeding

The study was conducted conforming to the local institutional laws and the Chinese law for the protection of animals. All adult zebrafish (Dario rerio) were maintained under standard conditions in accordance with our previous protocols [17, 18]. The *AB/WT*, *Tg(flk1:ras-mCherry)*, *Tg(kdrl:EGFP)*, *Tg(fli1a:EGFP)*, *Tg(fli1a:nEGFP)* and *Tg(mnx1:EGFP)* zebrafish used in this article has been described previously. Zebrafish embryos after 24 hpf were treated with 0.2 mM 1-phenyl-2-thio-urea to prevent pigment formation.

### Fluorescence-activated cell sorting

To compare the expression level of miR-22a in zebrafish neurons and ECs, *Tg(mnx1:EGFP)* and *Tg(Fli1a:EGFP)* larvae were used for neuron and ECs sorting, respectively. To investigate the effect of miR-22a sponge manipulations on the expression level of miR-22a in ECs and neurons, *Tg(mnx1:EGFP)* and *Tg(kdrl:ras-mCherry)* larvae microinjected *mnx1:EGFP-miR22a-S* and *flia:EGFP-miR22a-S* were used for both ECs and primary motor neuron sorting. Zebrafish embryos at 72 hpf were suspended by PBS containing 2% Fetal Bovine Serum (FBS) to remove the yolk. Then, embryos were digested using 0.25% trypsin at 28℃ for 30 min followed collecting by centrifugation. Collected cells were suspend with PBS (containing 2% FBS) and filtered with cell strainer (BD Falcon, 352340). The cells were used to sort ECs and neurons by a flow cytometer.

### RNA isolation, reverse transcription (RT), polymerase chain reaction (PCR), quantitative RT-PCR and RNA probe transcription

Total RNA of zebrafish embryos at various stages were extracted with TRizol according to the manufacturer’s instruction (Invitrogen, Waltham, MA, USA) and genomic contaminations were removed by DNaseI. Quantity of isolated RNA was verified using gel electrophoresis and Nanodrop, followed by cDNA synthesis using Transcriptor First Strand cDNA Synthesis Kit (Roche) then was stored at −20 °C. The custom-designed dre-miR-22a (Accession: MIMAT0001788) TaqMan® MicroRNA Assays were purchased from Thermo Fisher Scientific Inc. Assay ID : 004640_mat for dre-miR-22a, 000450 for hsa-miR-126, 002218 for hsa-miR-10b. The experiments were performed following the manufacturer’s protocol. TaqMan MicroRNA Reverse Transcription Kit (4366596) was used for the microRNA reverse transcription. Quantitative RT-PCR was conducted in a total 20 μl reaction volume with 10μl SYBR premix (TIANGEN). The relative RNA amounts were calculated with the comparative CT (2-DDCT) method and normalized with elongation factor 1-alpha (*ef1a*) as the reference. The primers for QRT-PCR are listed, for ef1a (Accession: NM_131263.1):

Forward primer: 5′ -TGA TCT ACA AAT GCG GTG GA-3′;
Reverse primer: 5′ -CAA TGG TGA TAC CAC GCT CA-3′.

Whole-mount in situ hybridization (WISH) with antisense RNA probes was synthesized as described previously [19]. The cDNA fragments used for miR-22a RNA probe transcription as template were amplified using the forward primer 5’-GAGGCCTCATCAGTTTGGAG-3’and reverse primer 5’-TCTCACTGCTCTG CATGCTT-3’. Digxigenin (DIG)-labeled sense and antisense probes were performed from the linearized pGEM-T-easy plasmids using the DIG RNA Labeling Kit (Roche).

### Whole-mount *in situ* hybridization

Zebrafish embryos were harvested at various stages, fixed overnight in 4% paraformaldehyde (PFA), washed with PBST, dehydrated in methanol then stored at 4 °C for subsequent use. The procedure for *in situ* hybridization follows our previous description [20]. For sectioning, above whole-mount embryos were transferred to Tissue-Tek OCT compound followed by being embedded in OCT blocks. The blocks were sectioned on a Leica RM2125 microtome at 10 μm.

### Injection of morpholinos (MO), microRNA precursor and construct

Morpholino antisense oligomers were synthesized according to the manufacturer’s protocol. The MO and miRNA precursor sequence is the following: Dre-miR-22a-MO: 5’-AGCTTGCCAGTGAAGAACTGCTGCA-3’; Dre-miR-22a control MO: 5’-CACAGATTCGGTTCTACTGCCTTAA-3’; rab11bb-MO: 5’-GCCATTTTAGACAAGCCGCCGCGTC-3’; miR-22a precursor :5’-GCUGACCUGCAGCAGUUCUUCACUGGCAAGCUUUAUGUCCUUGUGUACCAGCUAAAGCUGCCAGCUGAAGAACUGUUGUGGUUGGC-3’. Morpholinos were prepared and injected into single cell stage embryos as described previously. The *Tg(fli1a:EGFP-miR-22a-sponge)* and *Tg(mnx1:EGFP-miR-22a-sponge)* construct was injected into one cell stage *Tg(kdrl:ras-mcherry)*-fertilized egg (1 ng per embryo). In addition, the dCas9-KRAB (CRISPRi) approach was described in the published protocol [21].

### cDNA library preparation and sequencing

Total RNA was isolated and purified from 30 hpf zebrafish larvae using TRIzol Reagent (Invitrogen, USA). Then, RNA purity and integrity were verified by NanoDrop 2000 (Thermo Fisher, USA) and gel electrophoresis detection according to the protocol. A cDNA library was constructed using the TruSeq Ilumina RNA sample prep v2 kit by the manufacturer’s protocol. The final cDNA library was sequenced on an Illumina HiSeq 4000 (Illumina, San Diego, CA, USA).

### Identification of differentially expressed genes

Raw sequencing reads were assembled to the zebrafish reference transcriptome and genome (GRCz10 danRer10) using Bowtie2.0 and TopHat 2.0 (ref. 67). Differentially expressed genes (control vs miR-22-MO) were identified according to the value of Z2 fold using DESeq tool.

### Whole-embryo microRNA sensor assay in zebrafish

Whole-embryo microRNA sensor assay in zebrafish was carried out as described previously [20]. The coding sequences of EGFP and mCherry were cloned into the pCS2+vector. The pCS2+-EGFP-*sema4c*-3’-UTR construct was generated by cloning 1121 bp 3’-UTR of the zebrafishmib1Mrna (ENSDARG00000079611) into the pCS2+− EGFP vector, whereas pCS2+-EGFP-*sema4c*-3’-UTR (MUT) was generated by inserting only nucleotides 611 bp of the *sema4c* mRNA, which lacks the fragments containing the miR-10 targeting sites. *sema4c* 3’-UTR and mut *sema4c* 3’-UTR were inserted between the *EcoRI*-*XhoI* restriction sites in the multiple cloning regions downstream of the EGFP gene. The following two pairs of primers were used for cloning the insertion fragment:

*sema4c*-3’-UTR- EcoRI-left: 5’ -CCGGAATTCTGTGGTAGTTGAGGTGCTATCT -3’;
*sema4c*-3’-UTR-XhoI-right: 5’- CCGCTCGAGACAGTGTGAGCCAGCCTTAA -3’;
*sema4c* (mut)- 3’-UTR-EcoRI-left: 5’- CCGGAATTCTTGTGGTAGTTGAGGTGCTATC -3’;
*sema4c* (mut)- 3’-UTR-XhoI-right: 5’-CCGCTCGAGACTGGGCCTAATACACTATTGT-3’.

The pCS2+-mCherry vector was injection as a control. These three plasmids were linearized with Not1/Kpn1 and used as templates to synthesize the capped mRNAs using mMessage Machine (Ambion). The RNAs were injected into single cell stage embryos as described previously (35 pg per embryo).

### EC culture, oligos transfection, exosome isolation, and Imaging

HUVECs culture and oligos transfection experiments were performed as described previously [20, 22, 23]. The exosome isolation was described in our previous work [23]. For confocal imaging of nerves and blood vessel development in zebrafish, specific stage of larvae was anaesthetized and embedded in 0.6% low melting agarose. Confocal images were acquired with a Nikon TI2-E-A1RHD25 confocal microscope or a Leica TCS-SP5 LSM. Images were performed using Imaris software. For the results of in situ hybridization, images were taken by an Olympus stereomicroscope MVX10.

### Statistical analysis

Statistical analysis was performed by student’s t-test and one-way analysis of variance (ANOVA) using GraphPad Prism, in which p-values < 0.05 were considered statistically significant.

## Results

### miR-22a is highly expressed in endothelial cells in the zebrafish

To investigate the expression of miR-22a in endothelial cells of developing blood vessels, the EGFP+ cells in *Tg(kdrl:EGFP)* zebrafish embryos at 22 hpf were isolated by fluorescence-activated cell sorting (FACS) as previously described [20, 24]. Endothelial miRNA expression profiles were harvested according to deep sequencing as we previously described [20, 24] and miRNA quantitative PCR, showing that expression level of miR-22a were comparable to miR-126, which is generally accepted as a highly expressed miRNA in ECs (Supplementary Figure 1a, b). Furthermore, the expression profile of miR-22a in the zebrafish was analyzed using whole mount *in situ* hybridization (ISH) with a digoxigenin-labeled probe. In the developing embryos, miR-22a hybridization signal was detected in the blood vessels, which was consistent to the TaqMan miRNA assay and sequencing result (Supplementary Figure 1c).

### Deficiency of miR-22a caused aberrant vascular networks

Considering the highly expression of miR-22a in zebrafish embryonic ECs, it is rational to speculate it might modulate the development of blood vessel. To investigate the function of miR-22a during blood vessel development, miR-22a morpholino antisense oligonucleotide (miR-22a-MO) was injected into single cell stage of zebrafish embryos. This MO was designed to knockdown the expression of miR-22a through blocking Dicer and Drosha sites. The results of quantitative RT-PCR provided evidence that the injection of miR-22a-MO efficiently reduced the expression level of mature miR-22a (Supplementary Figure 2). The morphology of miR-22a deficiency was examined by confocal microscopy at different stages in blood vessel development. In contrast to control group, at 30hpf, the intersegmental vessels (ISVs) grew upwards halfway, then turned to horizontal sprouting (Figure 1a, b). In 48hpf, miR-22a-MO-injected embryos exhibited disorganized vascular networks (Figure 1a, c). Specifically, the patterning of disorganized vasculature can be classified into three types. The first case, the intersegmental vessels (ISV) in miR-22a knockdown zebrafish are not confined to one somite and connected to adjacent ISVs. The second, the ISV grew upwards halfway, then reversely extended to dorsal aorta. The third, the ISV grew halfway, then across somite to connect to opposite ISV or dorsal aorta. In addition, co-injection of miR-22a duplex significantly alleviated the disorganized ISV pattern, confirming the phenotype was a specific consequence of repression of miR-22a (Supplementary Figure 3a, b). Moreover, we generated a transgene *Tg(ubi:miR-22a-sponge)*, in which the mature miR-22a was competitively buffered to compromise function, and the ISVs displayed similar patterning defect with those of miR-22a morphants (Figure 1a). Taken together, these results suggest that miR-22a is required for the well-ordered pattern of vascular networks.

**Figure 1.**
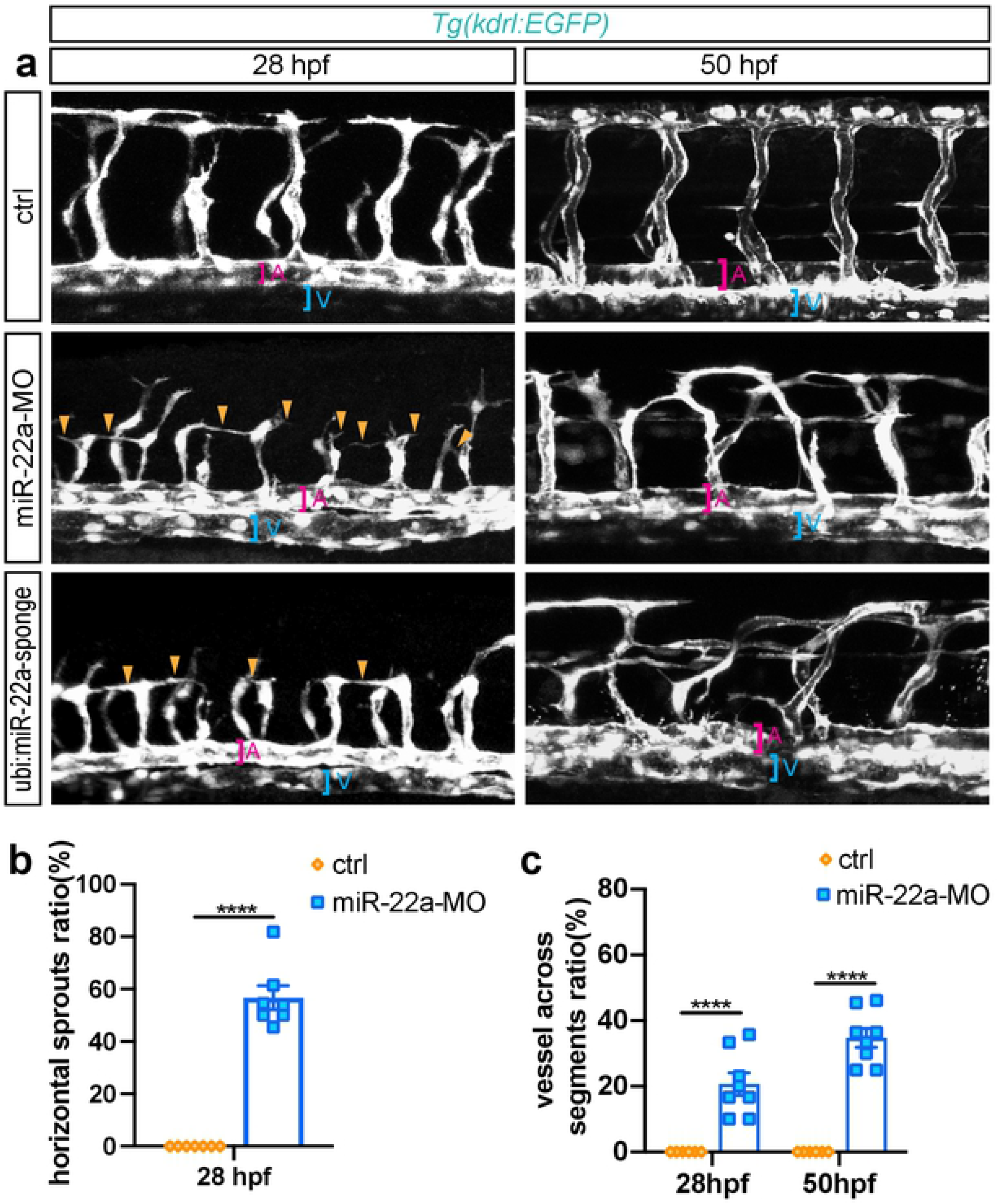
Deficiency of miR-22a caused aberrant vascular networks. (a) Confocal imaging analysis of control MO, miR-22a-MO and miR-22a-sponge-injected *Tg(kdrl:EGFP)* embryos at 30 hpf and 48 hpf. The arrowheads indicate the aberrant angiogenic sprouts. A: dorsal aorta; V: posterior cardinal vein. (b) The statistics of horizontal sprouts ratio in 30 hpf embryos injected with control MO, miR-22-MO and miR-22-sponge. t-test, ****, *p*<0.0001. (c) The statistics of embryos with vessel across segments ratio in each group: control MO, miR-22a-MO and miR-22-sponge. t-test, ****, *p*<0.0001.

### Knockdown of miR-22a impaired the pathfinding of tip cells

Endothelial tip cells have been proven to guide the proper wiring of nascent vessel though exploring the environment by filopodia. To further confirm the role of miR-22a in governing endothelial tip cell behaviors, time-lapse imaging were performed in *Tg(kdrl:EGFP)* zebrafish line. In the control, ISVs initiated to sprout from the dorsal aorta (DA) at 20 hpf, then grew upwards between somites and form dorsal lateral anastomotic vessel (DLAV) at around 30 hpf (Figure 2a, c; Movie 1). In the miR-22a deficiency embryos, ISVs emerged from DA at around 22 hpf and arrived at the horizontal myoseptum. Although the process of ISVs sprouting was normal, the subsequent migration of tip cells displayed multidirectional filopodial extensions and failed to reach the DLAV on the time (Figure 2b, d; Movie 2). Moreover, some tip cells stopped growing upwards and turned sideways or even backward extensions (Figure 2a, c). These results suggest that miR-22a seem not to be necessary for the initial stages of ISVs sprouting angiogenesis, but rather regulates the vascular patterning through affecting tip cell extension.

**Figure 2.**
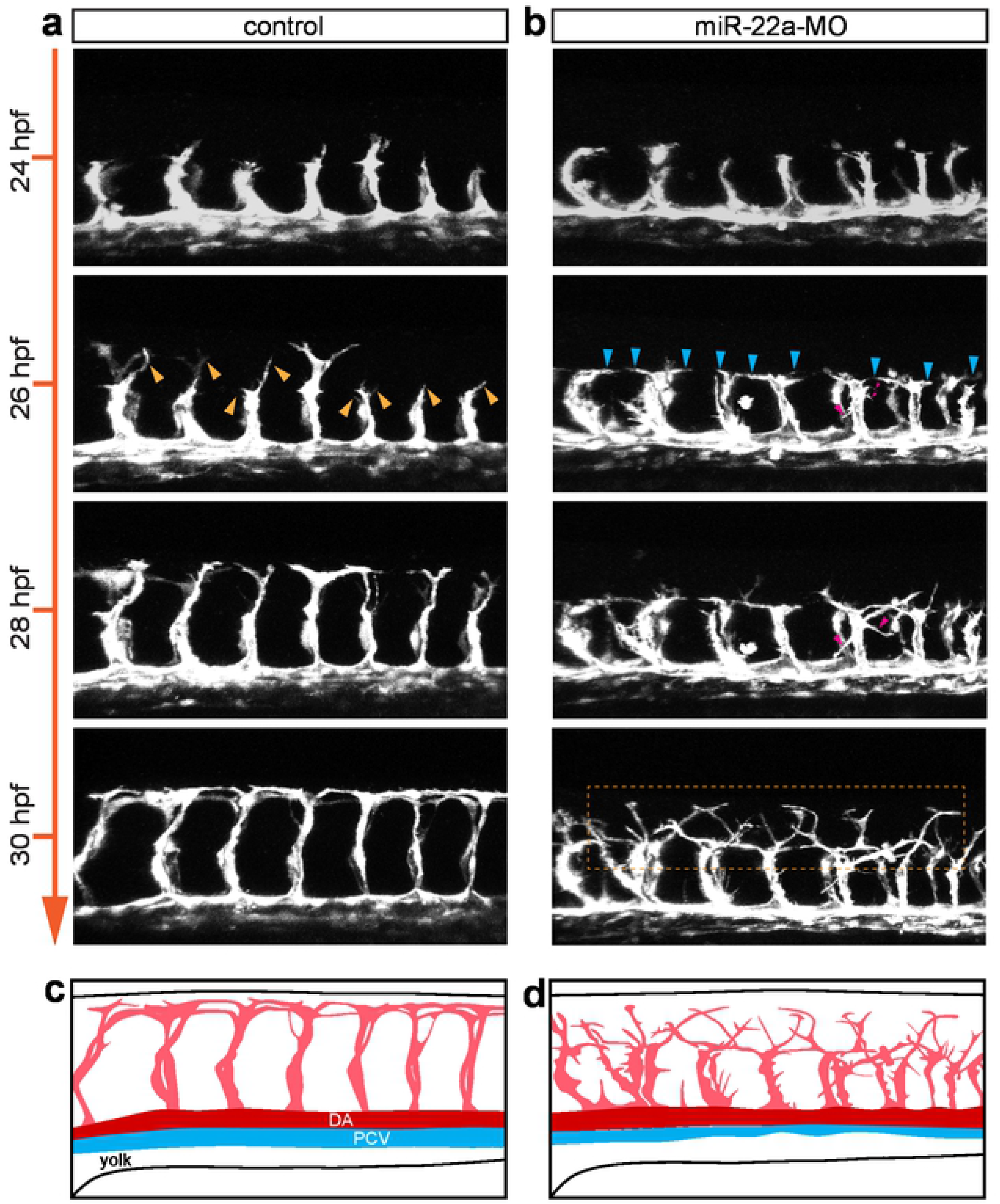
miR-22a regulates ISV tip cell behaviors. (a-b) Still images from in vivo time-lapse imaging analysis of *Tg(kdrl:EGFP)* embryos with HD detection setting. Time stages (hpf) are noted at the left side. (c-d) Diagram of ISV morphology in control and miR-22a MO-injected embryos.

### Deficiency of mir-22a caused aberrant axonal projection of PMNs

Since miR-22a was evidenced to express in neural system and required for ISV pathfinding, we reasoned that it might play a role in axonal projection of neurons. In order to investigate whether miR-22a regulates neuronal pathfinding, we examined the morphology of PMNs in the miR-22a knock down *Tg(mnx1:GFP)*^ml2^ zebrafish embryos at 48 and 72 hpf whose primary motor neurons were labeled with GFP using confocal microscopy imaging analysis. It was shown that absence of miR-22a caused dramatic developmental defects of PMNs (Figure 3a). Firstly, the axonal trajectories of Caps were significantly misled in the miR-22a morphants with nearly one third and half of Caps axons improperly across neighboring segments at 48 and 72 hpf respectively (Figure 3a-d). In contrast, it was barely to find in the control group (Figure 3a-d). Secondly, the deficiency of mir-22a induced the generation of the ectopic caps at 48 hpf (Figure 3a-c, e) and the ratio of this phenotype increased with the development of the zebrafish (Figure 3e). In order to confirm the motor neuronal defects were specifically caused by miR-22a inactivation, we carried out the confocal imaging analysis of the *Tg(mnx1:GFP::ubi:miR-22a-sponge)*, and the results revealed the similar motor neuron phenotypes with those in the miR-22a morphants, including the altered Cap trajectory and ectopic primary motor neuron (Figure 3a, b, d, e). Furthermore, we did the confocal time-lapse imaging analysis and found that the axonal trajectories of Caps exhibited the tendency to cross the somite at around 36 hpf (Supplementary Figure 4). From 48 hpf to 60 hpf, the axon of cap across the somite boundary continually extended parallel to spinal cord (Supplementary Figure 5).

**Figure 3.**
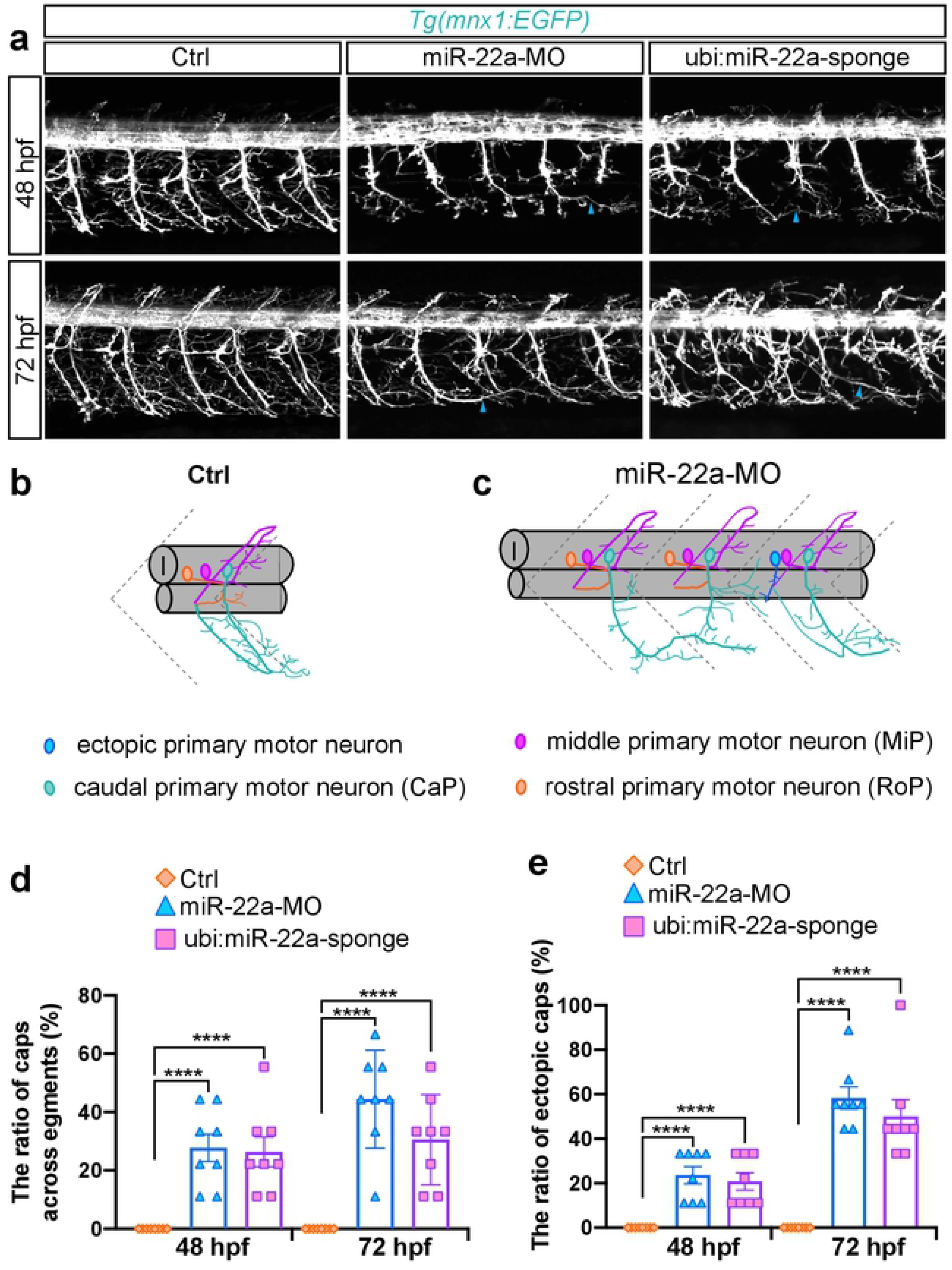
Deficiency of mir-22a caused aberrant axonal projection of PMNs. (a) Confocal imaging analysis of primary motor neurons in the control embryos, miR-22a morphants and *Tg(mnx1:GFP::ubi:miR-22a-sponge)* embryos at 48 and 72 hpf. Arrow heads indicate the caps across different segments. (b, c) The schematic diagram for primary motor neurons in the control embryos and miR-22a morphants. (d) The statistical analysis of the ratio of caps across different segments in the control, miR-22a morphants and *Tg(mnx1:GFP::ubi:miR-22a-sponge)* embryos at 48 and 72 hpf; One-way ANOVA, ****, *p*<0.0001. (e) The statistical analysis of the ratio of aberrant axonal projection of caps in the control, miR-22a morphants and *Tg(mnx1:GFP::ubi:miR-22a-sponge)* embryos at 48 and 72 hpf; One-way ANOVA, ****, *p*<0.0001.

### Endothelial miR-22a is required for both PMNs and ISV organization

To investigate the consequences of specific down-regulation of miR-22a expression level in ECs, we microinjected *Tg(fli1a:miR-22a-sponge)* constructs into one cell stage embryos, in which *miR-22a-sponge* was transiently expressed in ECs to block the function of miR-22a. The sponge contained seven repeats of the miR-22a antisense sequence, which inhibited miR-22a expression in ECs by chelating miR-22a. Comparing with the control group, embryos with ECs expressing *miR-22a-sponge* exhibited severe phenotypes of both PMNs and ISVs (Figure 4a, b), as we aforementioned in miR-22a morphants. Thus, ECs miR-22a is necessary for the pathfinding of ECs and neurons in zebrafish. We then explored whether miR-22a in neural system could regulate the pattern of nerves or vascular as well. To address this point, we specifically reduced the level of miR-22a in neurons though microinjection of *Tg(huc:miR-22a-sponge)*, in which *miR-22a-sponge* expression was driven by the promoter of the neuron-specific gene. In contrast to specific down-regulation of miR-22a in ECs, block the function of miR-22a specifically in neural system impaired the primary motor neuronal navigation, but less than 5% of those exhibited obvious vascular defects (Figure 4c, d). Taken together, these results suggested that endothelial miR-22a were involved in both vascular and neuron pathfinding, while neuronal miR-22a regulated neuronal navigation in cell-autonomous manner.

**Figure 4.**
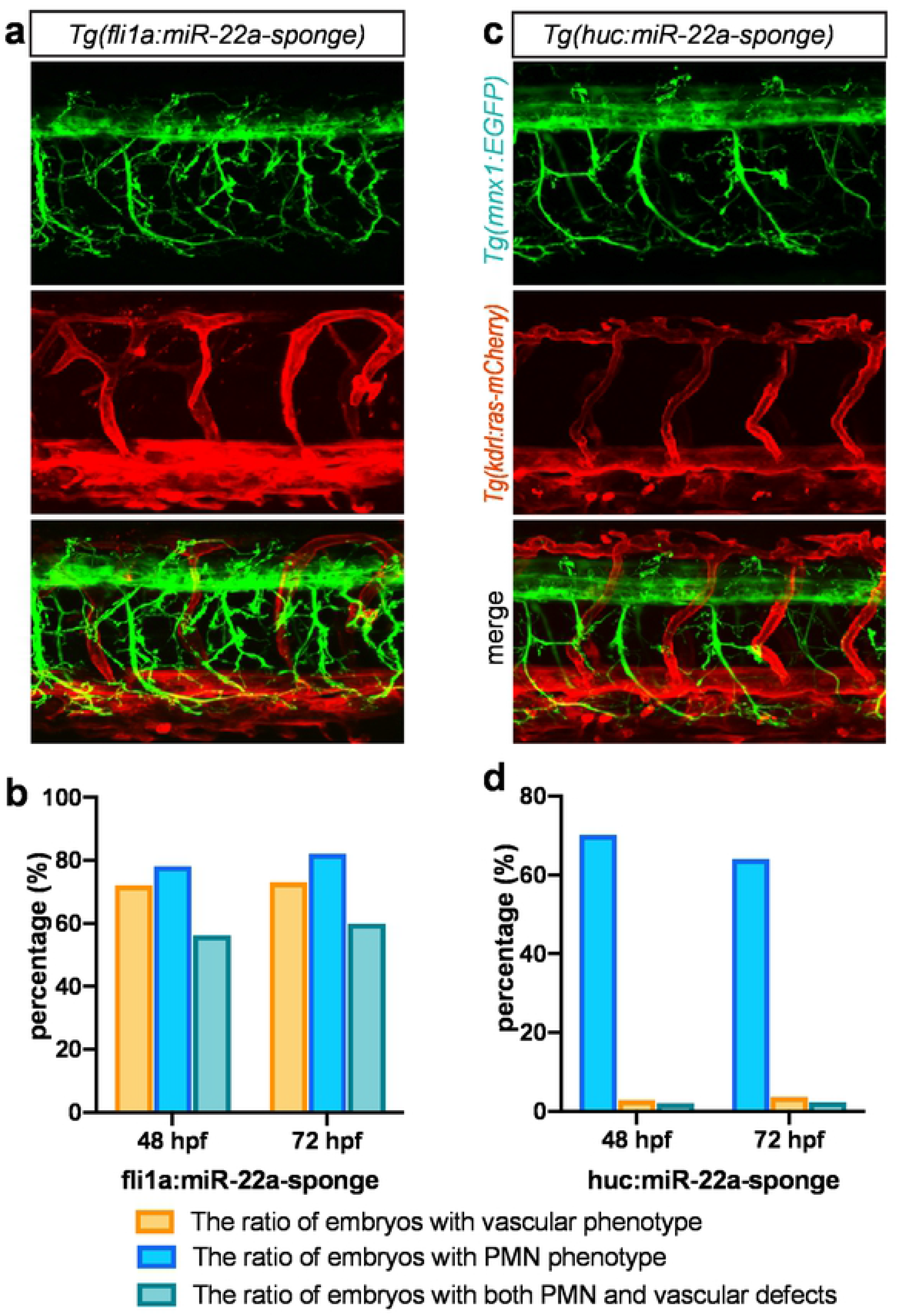
Endothelial miR-22a regulates primary motor neuron and ISV pathfinding. (a) Confocal imaging analysis of fli1a:miR-22a-sponge-injected *Tg(mnx1:EGFP::kdrl:ras-mCherry)* embryos at 72 hpf. (b) Percentage of embryos with impaired ISV and PMN pathfinding phenotype in fli1a:miR-22a-sponge-injected embryos at 48 hpf and 72 hpf, respectively. (c) Confocal imaging analysis of huc:miR-22a-sponge-injected *Tg(mnx1:EGFP::kdrl:ras-mCherry)* embryos at 72 hpf. (d) Percentage of embryos with indicated phenotypes in huc:miR-22a-sponge-injected embryos at 48 hpf and 72 hpf, respectively.

### Endothelial miR-22a regulates PMNs axonal navigation via exosome pathway

To explain why an endothelial-specific miR-22a knockdown embryo has defects in axons as well as blood vessels, and a neuron-specific miR-22a knockdown embryo also has axon pathfinding defects, we reasoned that the miR-22a of PMNs might be secreted by endothelial cells via exosome. The possible working model was illustrated in a diagram (Figure 5a). To verify this hypothesis, we carried out a serial of experiments. Firstly, we found that miR-22a was highly expressed in exosome isolated from human umbilical vein endothelial cells (HUVECs) by miRNA Taqman PCR analysis (Figure 5b). Furthermore, vesicles labeled with *Tg(fli1a:CD61-mCherry)* were detected in the zebrafish developing PMNs (Figure 5c), indicating endothelial derived exosome could be transported into PMNs. If the regulation of PMNs axonal navigation was involved in the exosome pathway, the blockage of the exosome formation would result in a phenotype of PMNs axonal navigation. To test this hypothesis, we inhibited the exosome generation through microinjection of the morpholinos targeting the blood vessel expressed Rab11(Rab11bb) and treatment of GW4869. It was found that block the generation of exosome resulted in the axon pathfinding defects of PMNs which were similar to the phenotypes caused by endothelial miR-22a loss of function (Figure 5e). In addition, we examined whether exosome isolated from endothelial cells could rescue the axonal navigation phenotype of PMNs in miR-22a morphants. We found that co-injection of the exosome isolated from cultured HUVECs partially rescued the axonal navigation defects (Figure 5f). Moreover, the percentage of phenotypes in embryos co-injected with exosome isolated from HUVECs, which were transfected with miR-22 duplex, was even less (Figure 5f). While, we further revealed that co-injection of the exosome isolated from HUVECs, which were transfected with miR-22 MO, failed to normalize the percentage of axonal navigation defects (Figure 5f). Taken together, these results indicated that the blood vessels modulated the primary motor neuronal pathfinding in zebrafish via exosome contained miR-22.

**Figure 5.**
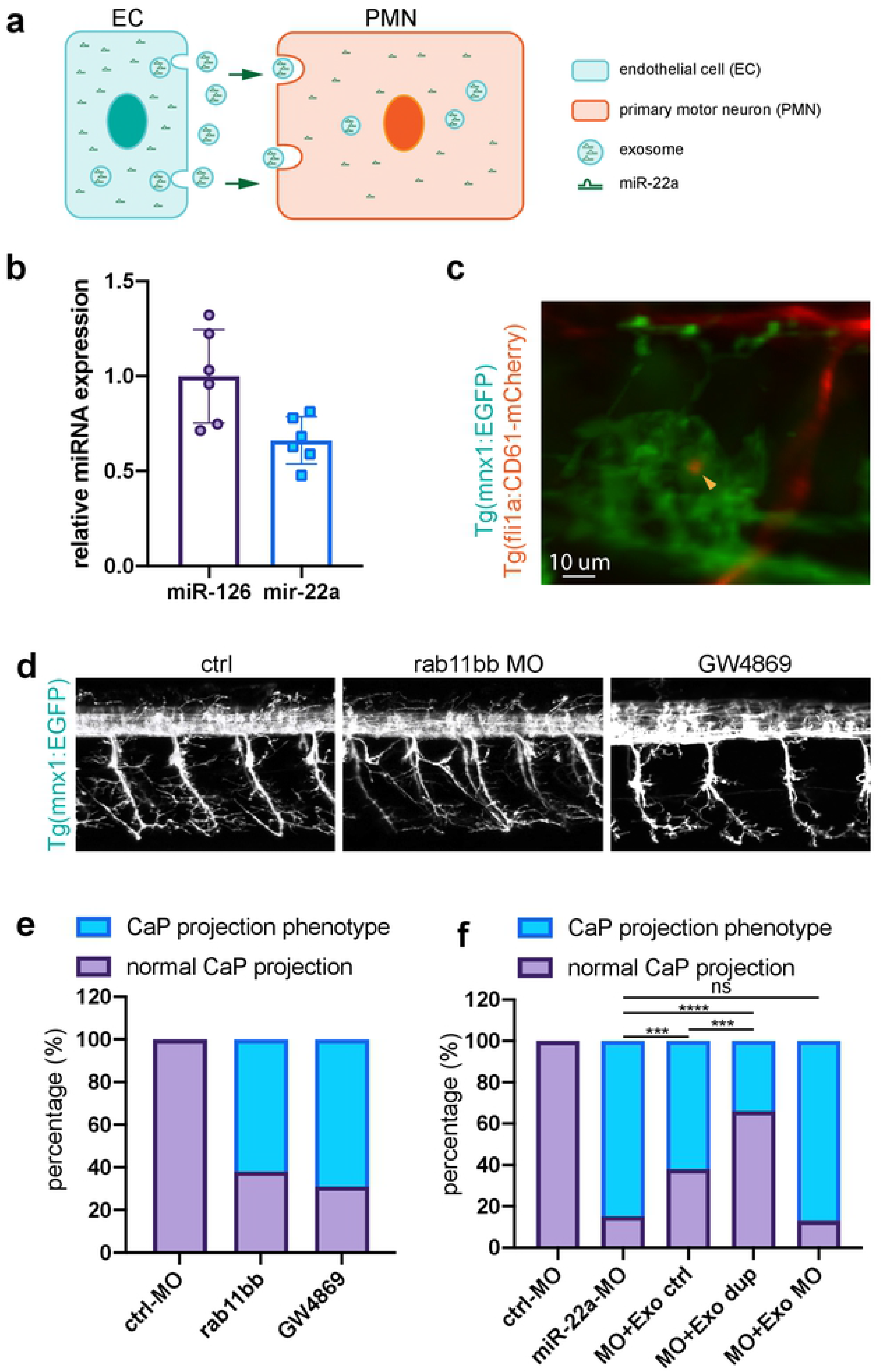
Endothelial miR-22a regulates PMNs axonal navigation via exosome pathway. (a) A working model for how blood vessels regulate primary motor neuronal pathfinding in zebrafish. (b) Q-PCR analysis of the miR-22 expression in the isolated exosome from HUVECs. (c) Imaging analysis of *Tg(mnx1:EGFP::fli1a:CD61-mCherry)*, arrowhead indicates the vesicle from endothelial cells. (d) Confocal imaging analysis of control, rab11bb MO injected and GW4869 treated *Tg(mnx1:EGFP)* embryos. (e) Percentage of embryos with indicated phenotypes in control, rab11bb MO injected and GW4869 treated *Tg(mnx1:EGFP)* embryos. (f) Percentage of embryos with indicated phenotypes in control, miR-22 MO injected, miR-22 MO co-injected with exosome isolated from HUVECs, miR-22 MO co-injected with exosome isolated from HUVECs transfected with miR-22 duplex, and miR-22 MO co-injected with exosome isolated from HUVECs transfected with miR-22 MO *Tg(mnx1:EGFP)* embryos. Fisher’s exact test, ****, *p*<0.0001; ***, *p*<0.001; ns, no significance.

### miR-22a targets *sema4c* in zebrafish embryos

In order to identify the direct target of miR-22a, we carried out a serial of experimental and *in silicon* analysis. The RNA samples from control and miR-22a knockdown embryo at 30 hpf for transcriptomic sequencing, which revealed 2191 up-regulated differentially expressed genes (DEGs) that might be affected by the repression of miR-22a. *In silico* analysis predicted that miR-22 potentially regulates thousands of genes in zebrafish using Targetscanfish (Release 6.2) and miRanda. A short list of overlapping target genes was selected from the transcriptomic DEGs and the predicted genes (Figure 6a). These potential target genes were selected for possible involvement in the regulation of vascular development and axon guidance including *kdr*, *sema4c*, *nppc*, *sema6d* and so on. Some class 4 semaphorins, such as *sema4a* and *sema4d* have been identified to participate in endothelial cell migration [10, 11]. Recently, *sema4c* and its receptor Plexin-B2 have been demonstrated to express in nervous system and endothelial cells, suggesting that *sema4c* may function as a guidance cue during endothelial cell development. *In silicon* analysis predicted that the 3’-UTR of *sema4c* in zebrafish contained potential miR-22a targeting site (Figure 6b). To confirm the functional interaction of miR-22a and *sema4c*-3’-UTR, the luciferase assay and EGFP sensor assay in zebrafish was performed, with the results indicated that the miR-22a precursor significantly inhibited the expression of *sema4c*-3’-UTR-EGFP but not *sema4c*-3’-UTR (mut), suggesting that miR-22a can directly target the zebrafish miR-22a-3’-UTR *in vivo* (Figure 6c, d). By quantitive PCR analysis, the mRNA expression levels of sema4c in ECs and PMNs in miR-22a injection embryos were significantly increased (Supplement Figure 6). These results suggest that miR-22a targets the 3’-UTR of *sema4c* and thereby guides endothelial cell behavior.

**Figure 6.**
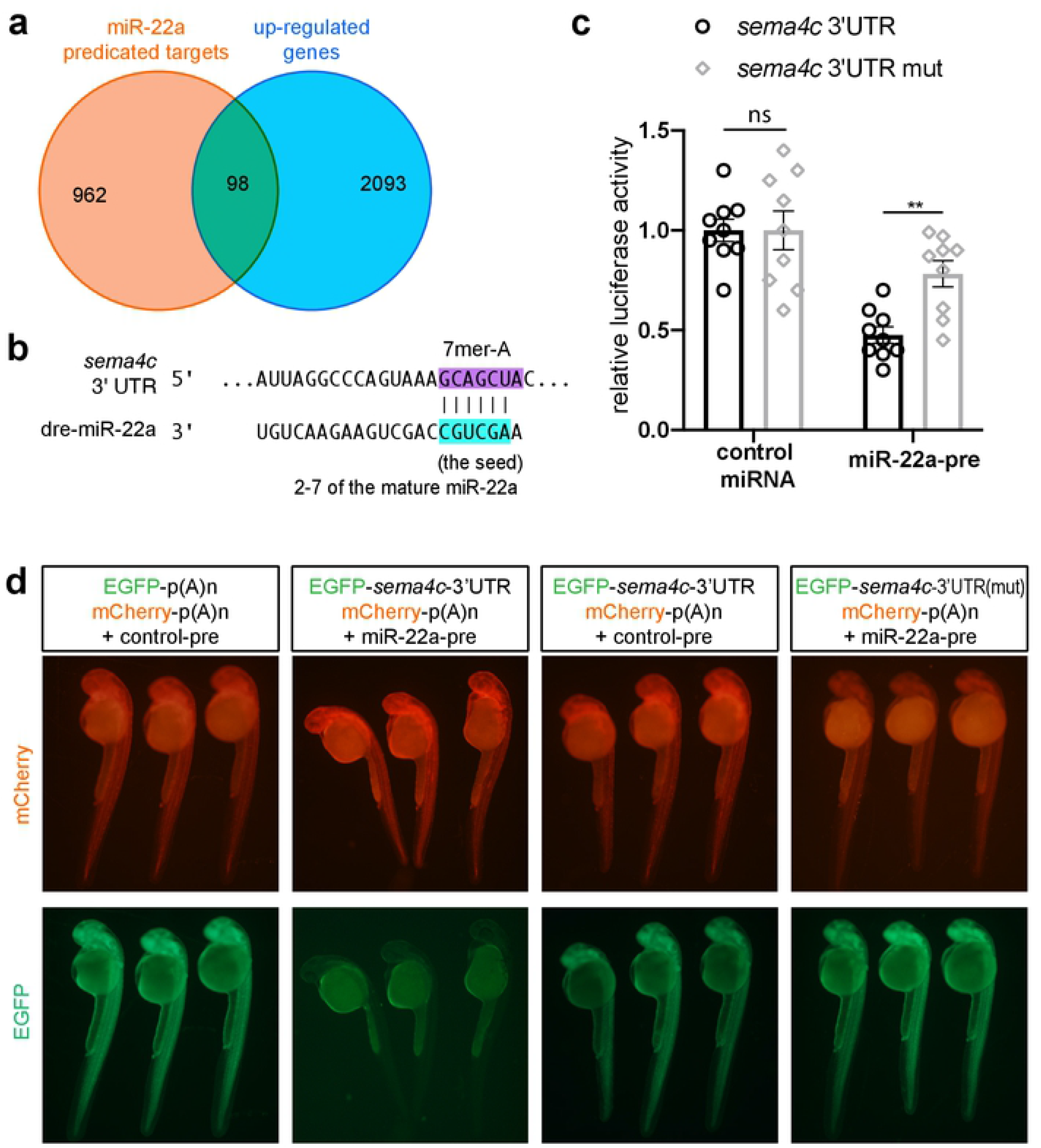
miR-22a directly targets *sema4c*. (a) Venn diagram of predicted target genes of miR-22a and transcriptomic up-regulated genes. (b) *Sema4c* 3’-UTR target sites of miR-22a. (c) Overexpression of miR-22 (miR-22-pre) reduced *sema4c*-3’-UTR luciferase activity in HeLa cells. Data are expressed as mean ± SE. t-test; **, *p*<0.01. (d) EGFP sensors were co-injected with mCherry control as indicated. miR-22a-precursor injection reduced the EGFP levels in EGFP-*sema4c*-3’-UTR sensor (second column), whereas mCherry levels were unchanged. In the mutated sensor, no reduction in GFP was noted (experiments were repeated three times; for each group, around 10 embryos were analyzed).

### Reducing *sema4c* partially restored the defects of ISVs and PMNs in miR-22a-deficient embryos

Our results suggested that up-regulation of *sema4c* was the likely cause of the defects of ISVs and PMNs in miR-22a-deficient embryos. If this was the case, the vascular and neuronal phenotypes of miR-22a-deficient embryos would be partially rescued by inhibiting the expression of *sema4c*. To investigate this possibility, *sema4c* was knocked down using dCas9-KRAB (CRISPRi) approach [21] in miR-22a morphants and then the ISV and primary motor neuron phenotypes were examined using confocal imaging analysis. Knockdown of *sema4c* in this setting greatly normalized both ISVs (Figure 7a, b) and PMNs projections (Figure 7c, d). This result suggests that miR-22a regulates vascular and primary motor neuronal pathfinding in zebrafish by directly targeting *sema4c*.

**Figure 7.**
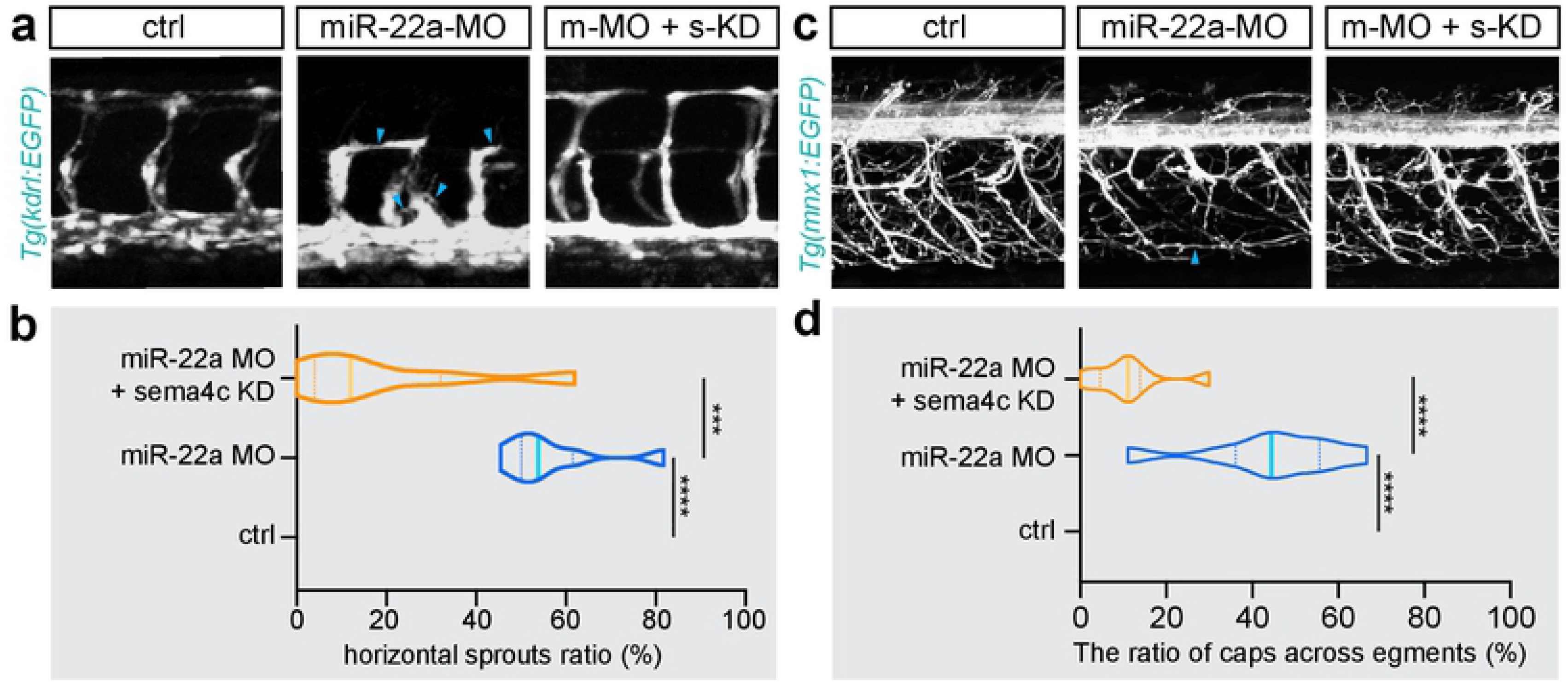
Reducing *sema4c* partially restored the defects of ISVs and PMNs in miR-22a-deficient embryos. (a) Confocal imaging analysis of blood vessels in control, miR-22a-MO, and miR-22a-MO + *sema4c* knockdown embryos at 30 hpf. (b) The statistics of horizontal sprouts ratio in control, miR-22a-MO, and miR-22a-MO + *sema4c* knockdown embryos at 30 hpf. One-way ANOVA; ***, *p*<0.001; ****, *p*<0.0001. (c) Confocal imaging analysis of control, miR-22a-MO, and miR-22a-MO + *sema4c* knockdown *Tg(mnx1:EGFP)* embryos at 72 hpf. (d) The statistical analysis of the ratio of aberrant axonal projection of caps in the control, miR-22a morphants and miR-22a-MO + *sema4c* knockdown embryos at 72 hpf; One-way ANOVA, ****, *p*<0.0001.

## Discussion

In this study, detailed expression analysis showed that miR-22a was highly enriched in endothelial cell and neural system of zebrafish embryos. Morpholino knock-down of miR-22a perturbed the pattern of ISVs and PMNs, indicating a requirement for miR-22a in directing endothelial and neuronal navigation during zebrafish embryonic development. Since the neuronal block of miR-22a specifically caused the axon pathfinding defects of PMNs, we excluded that the emergence of aberrant axonal navigation of PMNs was due to a consequence of vascular defects. Furthermore, we revealed that endothelial miR-22a regulates PMNs axonal navigation via exosome pathway. Target prediction analysis indicated that miR-22a potentially regulates thousands of genes. With transcriptomic analysis, 2191 up-regulated differentially expressed gene (DEGs) that might be affected by the repression of miR-22a were identified. A short list of overlapping target genes was selected from these DEGs and predicted genes. Combined with the sensor assay in zebrafish, *sema4c* was identified as a potential target. Then, the levels of *sema4c* were found to be elevated in miR-22a morphants by quantitively PCR analysis. In addition, inhibition of *sema4c* partially rescued the vascular and neuronal pattern in miR-22a morphants, suggesting that the observed phenotypes were caused by up-regulation of *sema4c*.

The vascular system is so complicated that different cell types and signal factors are coordinated to regulate the development processes. In particularly, the precise wiring of the vascular network is regulated by several guidance cues that dynamically modulate the endothelial cell-to-cell behavior. Currently, accumulating evidences have indicated that the guidance of blood vessel organization shares the similar or common cues involved in nerves network, which share multiple parallelisms of the morphological and functional feature with vascular system. ECs can respond to multiple axon-guidance factors during vascular development. Conversely, vascular guidance signal may also regulate neuronal development [25]. For example, the work of Calvo et al., indicated that VEGFR3 signaling can influence neurogenesis as well as angiogenesis [26]. Besides, vessels can also produce axon-attracting cues for neurons development [27, 28]. Sema4c is a member of class 4 semaphorins, which are a large family of secreted and membrane-bound cues that has key roles in vascular pattering and axon guidance [29]. Sema4c has long been considered associating with neuronal migration, but recent report found that Sema4c and its receptor Plexin-B2 were also expressed in endothelial cells [15]. In addition, Sema4c was reported to promote angiogenesis in breast cancer [30]. Although Sema4c is a transmembrane protein, its extracellular domain near the transmembrane portion can be cleaved [31]. By far, a few of miRNAs have been reported to participate in blood vessel formation by targeting the components of semaphoring family [32]. Here we describe an endothelial-enriched miRNA, miR-22a, which plays a crucial role in zebrafish vascular and neuronal pattern via regulation of *sema4c*. Loss of function of miR-22a in zebrafish embryo led to aberrant patterning of ISV and primary motor neurons, suggesting that miR-22a regulated ECs and neurons pathfinding. As the miR-22a expressed at a high level in the endothelial cells, it is possible that the phenotypes we have identified were attributed to the miR-22a from endothelial cells. We further investigated whether endothelial-derived miR-22a make dual roles in vascular and nerves system, the consequences of specific down-regulation of miR-22a in endothelial cell and neuron cell were compared, respectively. It was found that all these phenotypes can be observed by inhibition of miR-22a in endothelial cells. However, only aberrant neuronal pattern was phenocopied by loss of miR-22a in neuron cells. These data support the hypothesis that endothelial-derived miR-22a in zebrafish functions as a guidance signal regulates both vascular and neuronal branching and pathfinding. In addition, neuron-derived miR-22a could also regulate its patterning, while exert no significant influence on vascular network. We provided evidences to show that endothelial miR-22a regulates PMNs axonal navigation via exosome pathway. These results explained why an endothelial-specific miR-22a knockdown embryo has defects in axons as well as blood vessels, and a neuron-specific miR-22a knockdown embryo also has axon pathfinding defects.

To date, limited studies have reported the discovery and functions of miRNAs in vascular or nerves patterning [33]. Accumulated reports suggested that ECs and neurons can recruit various common guidance factors to control patterning of the complex vascular and neuronal networks. However, studies regarding the functional overlap between vascular and neuronal pathways, especially how the vessel contributes to neurovascular co-patterning is still in its infancy. In present study, we identified miR-22a, whose target gene *sema4c* is not only expressed in ECs, but also in neurons. Furthermore, miR-22a was revealed to play a crucial role in zebrafish blood vessel and neuronal patterning. Taken together, our findings provide a notable case that an endothelial-derived miRNA is attributed for regulating both ECs and neuronal pathfinding.

## Acknowledgements

This study was supported by grants from the National Natural Science Foundation of China (2018YFA0801004 and 81870359 received by Dong Liu; 81970432 received by Yunwei Shi; http://www.nsfc.gov.cn), Natural Science Foundation of Jiangsu Province (BK20180048, and BRA2019278 received by Dong Liu; http://kjjh.jspc.org.cn).

## Author Contributions

DL supervised and designed this project. JS, DL, JG wrote the manuscript. DL, JS, and JG analyzed the data. JS, JG, YS and XW performed the experiments.

## Supplementary Figure Legends

**Supplement Figure 1. Endothelial expression of miR-22a in zebrafish.** (a) The relative expression of miR-22a compared with miR-126, which are a highly expressed miRNA in ECs. (b) The relative expression of miR-22a compared with miR10s and miR-126s using Taqman PCR analysis. (c) Whole-mount in situ hybridization analysis of miR-22a expression in the zebrafish embryo at 30 hpf. Arrow heads indicate blood vessels.

**Supplement Figure 2. miR-22a-MO injection efficiently reduced levels of mature miR-22a of zebrafish embryo.** Quantitative PCR analysis of mature miR-22a in control-MO, miR-22a-MO embryos at 24 hpf, 48 hpf, and 72 hpf. t-test; ****, *p*<0.0001.

**Supplement Figure 3. Co-injection of miR-22a-MO and miR-22a duplex significantly alleviated the disorganized ISV pattern.** (a) Confocal imaging analysis of control, miR-22a-MO, and miR-22a MO and duplex mixture injected *Tg(kdrl:EGFP)* embryos at 48 hpf. (b) The statistics of caps across segments ratio in embryos injected with control, miR-22a-MO, and miR-22a MO and duplex mixture. One-way ANOVA; ****, *p*<0.0001.

**Supplement Figure 4. Deficiency of miR-22a caused aberrant axonal trajectories of caps.** (a) Diagram illustrating the imaging position. (b) Still images from in vivo time-lapse imaging analysis of axonal trajectories of caps in *Tg(mnx1:GFP)* embryos from 24 to 36 hpf. Arrowheads indicate the aberrant axon.

**Supplement Figure 5. Deficiency of miR-22a caused the parallel extension of caps.** (a) Diagram illustrating the imaging position. (b) Still images from in vivo time-lapse imaging analysis of axonal extension of caps in control and miR-22-MO-injected *Tg(mnx1:GFP)* embryos from 48 to 60 hpf. Arrowheads indicate the aberrant axon.

**Supplement Figure 6. miR-22a-MO injection efficiently increased levels of *sema4c* in ECs and PMNs.** (a) Quantitative PCR analysis of sema4c in control ECs and miR-22a-MO-injected ECs. t-test; **, *p*<0.01. (b) Quantitative PCR analysis of sema4c in control PMNs and miR-22a-MO-injected PMNs. t-test; ****, *p*<0.0001.

---

**Movie 1. Confocal time-lapse imaging analysis of intersegmental vessel formation in control *Tg(kdrl:EGFP)* embryos.**

**Movie 2. Confocal time-lapse imaging analysis of intersegmental vessel formation in miR-22a MO-injected *Tg(kdrl:EGFP)* embryos.**

